# Assessing the impact of temperament on training efficiency in *ex-situ* Red Brocket Deer (*Mazama americana*)

**DOI:** 10.1101/2024.11.28.625851

**Authors:** Lara Caveanha Gragnanello, Mariana Parra Cerezo, Cristiane Schilbach Pizzutto, Mateus José Rodrigues Paranhos da Costa

**Author notes:** Corresponding authors (LCG).

## Abstract

The personality of wild animals has been extensively discussed due to its significant influence on *ex-situ* management and conservation methods. However, there is still limited research directly addressing its impact on habituation and conditioning processes, which are essential techniques in conservation centers, zoos, and research institutions. This study aimed to demonstrate how individual personality variation affects training. We conducted the research at the Deer Research and Conservation Center of the Faculty of Agricultural and Veterinary Sciences, São Paulo State University, in Jaboticabal-SP, Brazil, with twelve red brocket deer (*Mazama americana*). Temperament tests were performed at the beginning of the study, assessing the animals’ reactivity to the presence of an unfamiliar human and during handling procedures. Following these tests, we initiated the processes of habituation and training of the animals. Three deer did not habituate to the trainer’s presence, being not subjected to operant conditioning; and four did it in just seven sessions. Among the nine deer that participated in the conditioning process, only five learned all the commands, representing a success rate of 41.66%. We concluded that more reactive animals, characterized as nervous and agitated, exhibited poorer performance in training. In contrast, less reactive animals, which were calmer and either allowed to approach or approached voluntarily an unfamiliar person, learned the commands more effectively.

## 1. Introduction

Temperament, often synonymous with personality, refers to the unique set of characteristics that determine consistent patterns of feelings, thoughts, and behaviors over time and across different conditions within individuals of the same species [1–3]. Understanding temperament is essential for grasping the interactions between individuals and their environment [4], particularly when implementing appropriate management practices and promoting animal welfare [5]. Research has shown that temperament can influence learning capacity [6] and responses to human interactions [7,8].

In cervids, research on temperament is particularly relevant due to their reactive and wary behavioral traits, essential for survival in the wild [9,10]. However, when under human care, these inherent behaviors complicate veterinary management and impede early disease identification, posing significant challenges to cervids conservation under human care [11], and this is particularly relevant when considering *ex-situ* management of red brocket deer (*Mazama americana*). This species can reach up to 65 cm in height and 145 cm in length and weigh between 30 and 40 kg [12]. According to the IUCN classification, *M. americana* is categorized as “Data Deficient” [13], underscoring the importance of studying its temperament and conditioning to aid conservation efforts.

According to Duarte et al. [14], physical restraint is discouraged because of the power of the hind legs, which poses a risk of injury to personnel. Consequently, chemical restraint often becomes the only viable option. Although this method can facilitate handling, it induces physiological changes in the animals and can affect their behavior for several days [4], further complicating their management.

Considering these challenges, operant conditioning training emerges as a valuable tool in conservation centers and zoological institutions [15]. This learning process shapes behaviors through positive reinforcement, establishing associations between specific actions and their outcomes [16]. Operant conditioning training is crucial for developing management routines, as it encourages animals to respond to commands and interact positively with caregivers and veterinarians [17,18]. By utilizing positive reinforcement, this approach not only significantly enhances the quality of life of the animals but also minimizes the necessity for physical and chemical restraints during management [19].

Despite some advancements, gaps still exist in understanding how the individual temperamental characteristics of deer influence their responses to conditioning. Furthermore, the relationship between temperament and learning is underexplored in this species, leaving room for deeper investigations. This study aimed to clarify how differences in temperament impact the success and effectiveness of training of Red Brocket deer. The results of this study have the potential to inform more effective management practices, enhance the quality of life for animals’ *ex-situ*, and contribute to conservation strategies that recognize the importance of temperament in adaptation and survival.

## 2. Material and methods

This study was approved by the Ethics Committee on the Use of Animals of the Faculty of Agricultural and Veterinary Sciences, UNESP, Jaboticabal, SP, Brazil (protocol number 710/21).

### 2.1 Animals and location

The study was conducted with 12 red brocket deer (*Mazama americana*) at the Deer Research and Conservation Center (NUPECCE, registered in SisGen, number AC0AA2A) of the Faculty of Agricultural and Veterinary Sciences at São Paulo State University, in Jaboticabal-SP, Brazil. The subjects included both males and females of various ages, all born and raised at NUPECCE. Their characterization is detailed in Table 1.

**Table 1.**
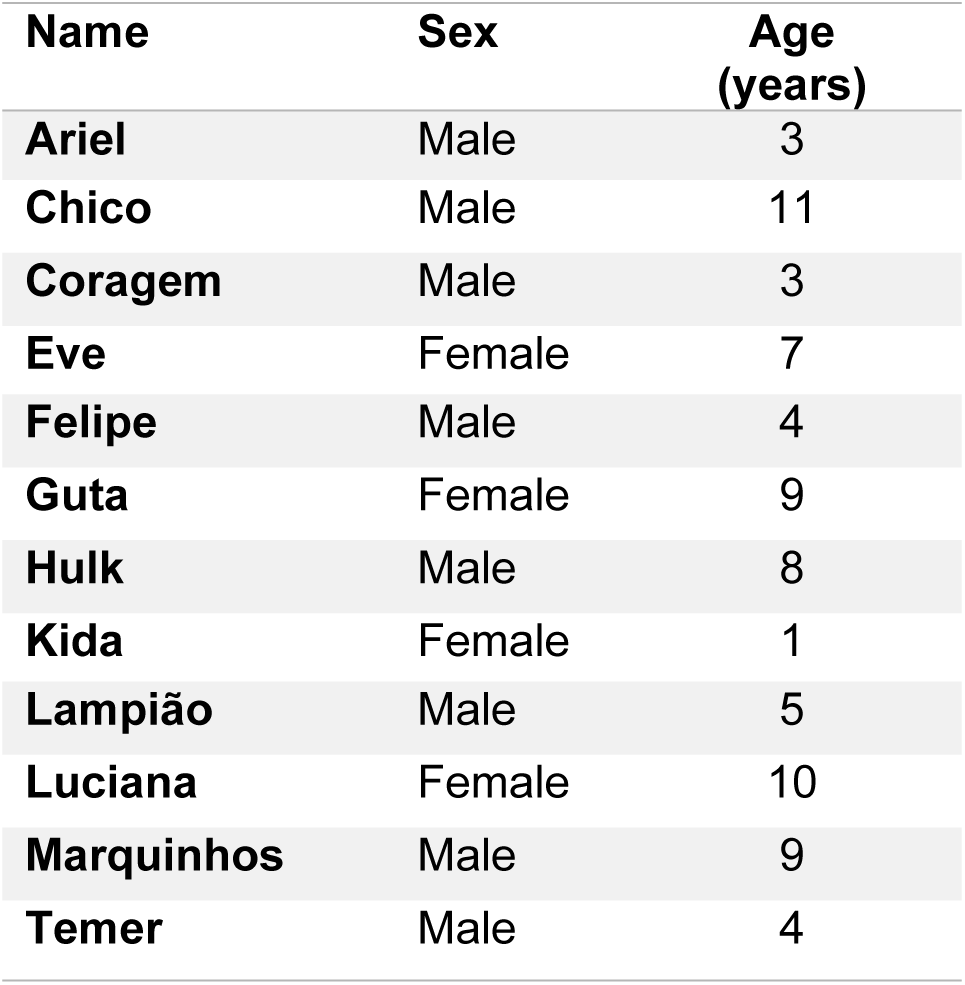
Name, sex, and age of the subjects of this study.

| Name | Sex | Age (years) |
| --- | --- | --- |
| Ariel | Male | 3 |
| Chico | Male | 11 |
| Coragem | Male | 3 |
| Eve | Female | 7 |
| Felipe | Male | 4 |
| Guta | Female | 9 |
| Hulk | Male | 8 |
| Kida | Female | 1 |
| Lampião | Male | 5 |
| Luciana | Female | 10 |
| Marquinhos | Male | 9 |
| Temer | Male | 4 |

### 2.2 Temperament assessment

Four temperament tests were conducted before the learning processes to define the individuality of the deer. The tests were carried out by the same two individuals, with one having no prior contact with the animals (unfamiliar), considering three different contexts: in the stall (reactivity in the presence of an unfamiliar person), during the transfer to the handling box (reactivity when moved to the handling box), and in the handling box (reactivity when inside the handling box and reactivity when touched while restraint in the handling box), as described below.

#### 2.2.1 Reactivity test in the presence of an unfamiliar person

This test was conducted in accordance with the methodology established by Waiblinger et al., [20], with necessary adaptations to suit the study’s framework. An unfamiliar person slowly opened the stall door and positioned herself so the animal could see her. The unfamiliar person remained at the door for 2 minutes without making any moviments, allowing the deer to habituate to her presence. In the following 10 minutes, it was recorded whether the animal approached the unfamiliar person (1) or not (0). After 12 minutes, the unfamiliar person slowly approached the deer, extending their right arm to try to touch it on the neck, and the animals were scored based on their reactivity, as described in Table 2.

**Table 1.** Descriptions of the reactivity scores assigned to deer *(M. americana*) when approached by an unfamiliar person.

| Score | Reactivity | Description |
| --- | --- | --- |
| 1 | Extremely calm | Deer allows being touched by an unfamiliar individual without attempting to escape. |
| 2 | Calm | Deer remains calm but does not allow being touched, moves around the enclosure, and allows being touched but shows some reluctance. |
| 3 | Slightly nervous | Deer shows moderate agitation and avoids the approach of the unfamiliar person but does not exhibit aggressive behavior or sudden movement. |
| 4 | Nervous | Deer runs back and forth in the enclosure when approached by the unfamiliar person but does not attempt to jump over the walls or hit the ground with its front paw to escape. |
| 5 | Extremely nervous | Deer attacks the unfamiliar person or runs back and forth in the stall, frequently looking up and attempting to jump against the walls. |

#### 2.2.2 Reactivity tests during handling procedures

On a different day from the tests previously described, we evaluated the reactivity of the deer when transferring them from the stall to the handling box. During this test, we recorded whether the deer jumped (over 1 meter high) or not (1 = yes or 0 = no) while being driven from the stalls to the handling box. The handling box measures 108 cm x 82 cm x 38 cm and features two guillotine-style gates on the shorter sides that open by pulling the lid upwards. Additionally, it has eight holes on the sides and two guillotine-style windows on the top (Fig 1).

**Fig 1.**
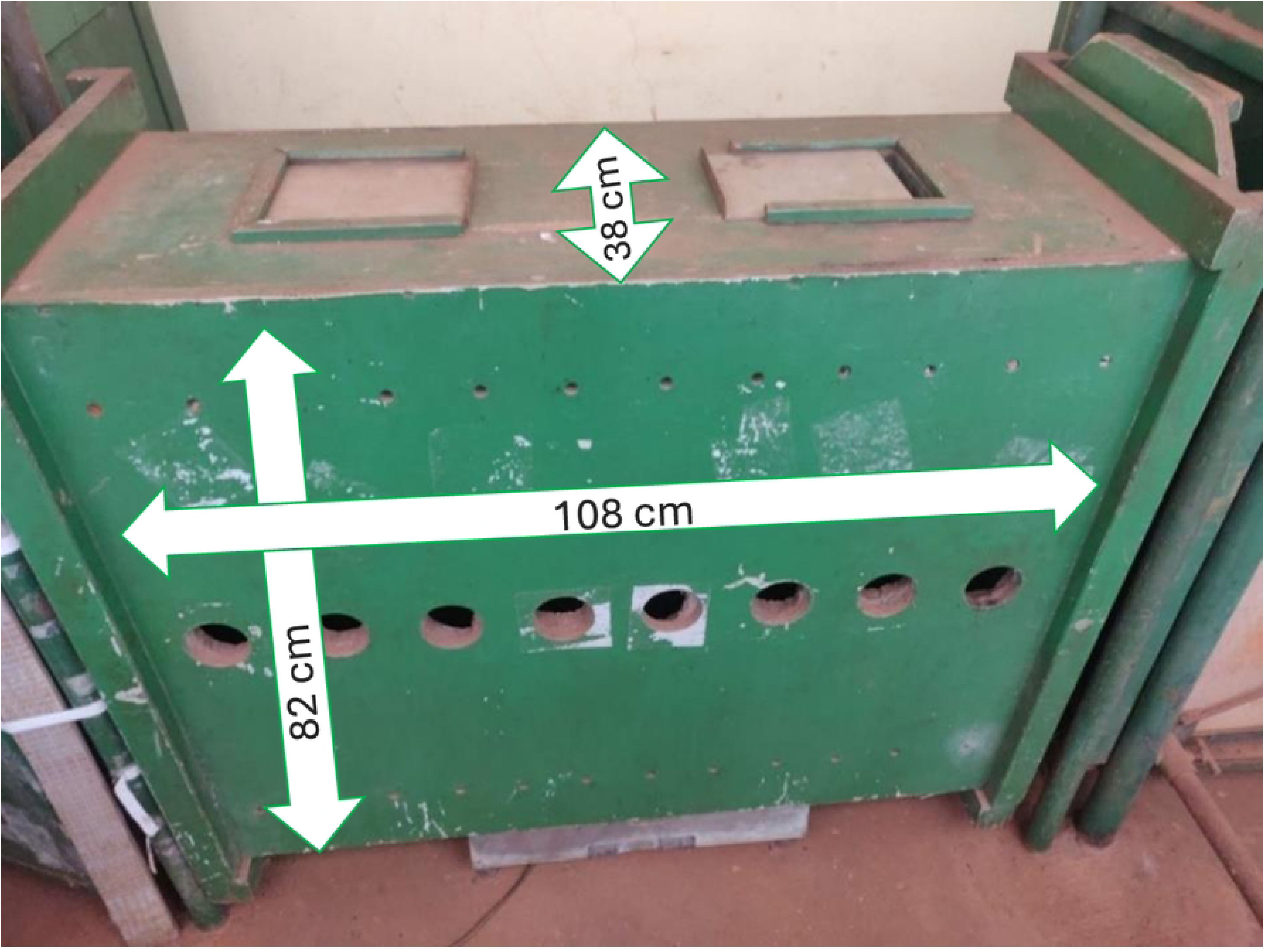
Dimensions of the handling box used for physically restraining red brocket deer (*M. americana*) during veterinary procedures and temperament tests for this study.

After each deer entered the box, the gates were closed, and the following tests were conducted. While the deer was restrained in the handling box, the trainer opened one of the upper windows and gently touched its hindquarters. Based on their responses, the trainer assigned one of the following movement scores:

1. No movement.
2. Minimal movement, lacking vigor, and remaining stationary for more than half of the interaction time.
3. Minimal yet vigorous movement.
4. Frequent and vigorous movement, characterized by back-and-forth motion within the box.
5. Excessive movement, including jumping and attempts to escape by pushing against the box with its hindquarters.

Following the trainer’s touch on the backs of the deer, the body posture was scored as (1) when standing, (2) when kneeling, or (3) when lying in a sternal position.

### 2.3 Learning processes

The training process was divided into two phases. The first phase focused on habituating the deer to the trainer and fostering positive interactions (S1A Fig). In the second phase, operant conditioning techniques were applied to train the deer for veterinary management practices. Throughout the learning process, the materials used were a *target* to guide the desired movements and actions, primary rewards (pieces of banana), and secondary rewards that included verbal praise (e.g., ‘very good’) and a clicker to reinforce positive behaviors.

Each habituation session lasted approximately 10 minutes, beginning with the trainer opening the stall door. Throughout each session, the trainer interacted with the deer by calling them by name while positioning a chair and a food container with feed in place. Holding a bowl of banana pieces, the trainer maintained continuous verbal interaction with the deer. After roughly eight minutes, she offered the banana pieces to the deer. If the deer approached, she extended the bananas by hand; otherwise, he would toss them nearby. This procedure continued until the deer displayed calm behavior or reduced escape responses. Over time, the trainer began to enter the stall, moving slowly among the deer, calling to it, and showing the banana pieces.

The second phase focused on operant conditioning (S1B Fig), grounded in continuous reinforcement, whereby each correct command received a reward. This phase commenced with the deer calmly approaching the trainer as she walked through the stall. Training sessions took place daily for each deer, incorporating at least one rest day weekly, occurring between the conclusion of the first management session at 9:00 AM and the initiation of the second at 2:00 PM. Furthermore, all sessions underwent recording for documentation and analytical purposes.

Before opening the stall, the trainer repeated the deer’s name three times. Upon entering, she offered a piece of banana. If the deer did not approach, she would bring the fruit to it. Subsequently, she positioned herself at a corner of the stall and initiated the conditioning process with a familiar command “come”.

The trainer utilized a trace-conditioning method: upon giving a command and receiving a correct response from the deer, she pressed the clicker before offering pieces of fruit as a reward. Training success involved the deer approaching after hearing the voice command without seeing the reward. At the end of each session, a container of feed placed inside the enclosure received a topping piece of banana.

The operant conditioning commands were shown according to the individual progress of the deer. For example, deer that reached the command “belly” had already successfully performed the commands “come” and “snout.” Operant conditioning always began with commands that were already learned and was always concluded with the command “over”, during which a larger quantity of fruit was provided. The information about the commands applied during training is detailed in Table 3.

**Table 3.**
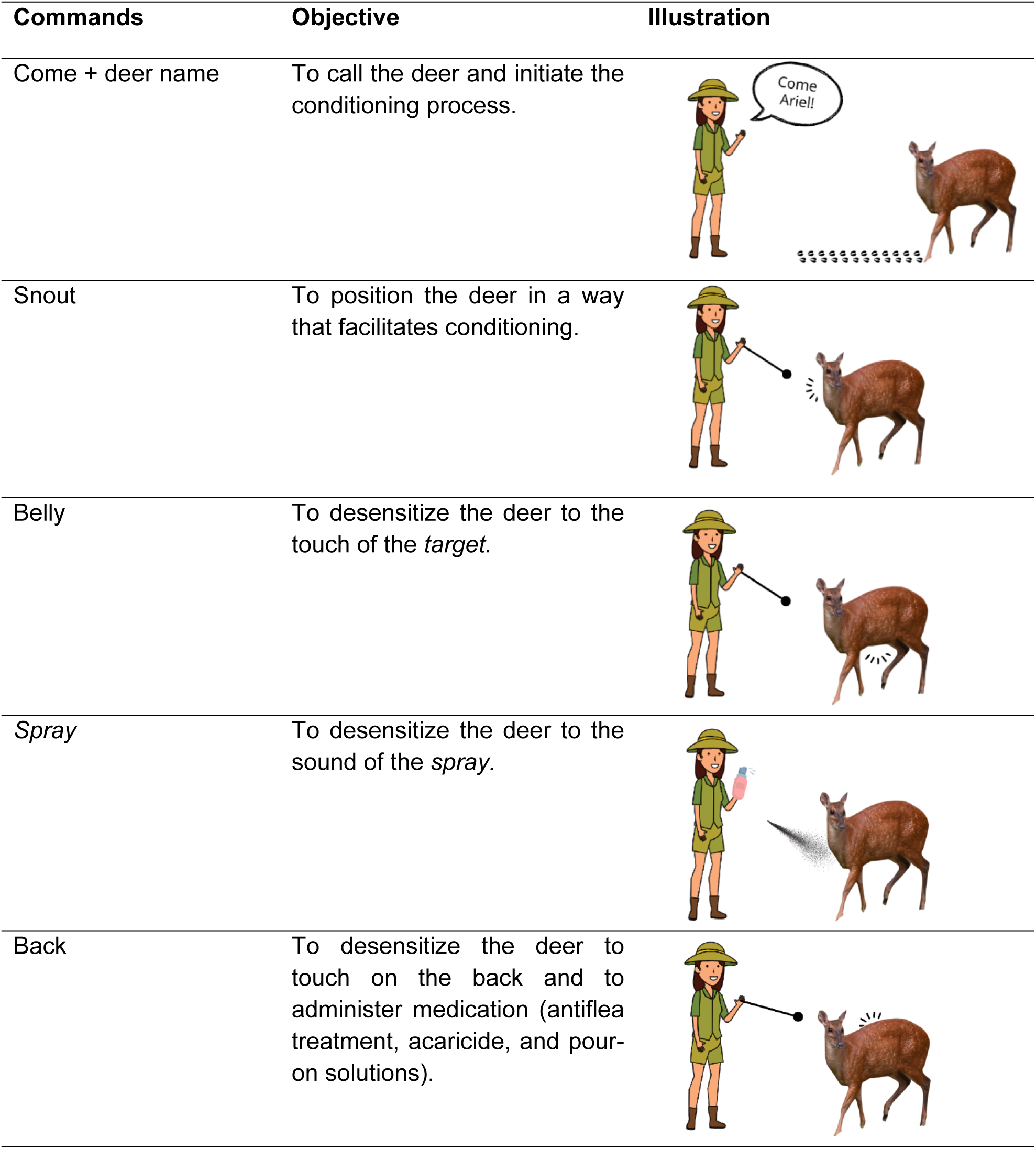
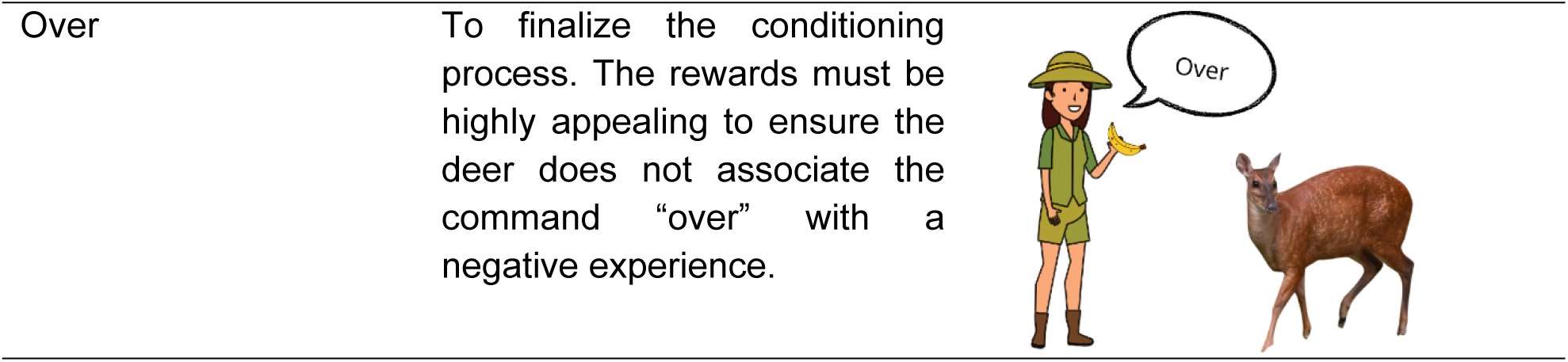
Commands applied during operant conditioning, respective objectives, and illustrations displaying the practical application of each command.

### 2.4 Statistical analysis

The statistical analyses were conducted using R software, version 4.3.3. A multiple correspondence analysis (MCA) was applied to reduce the categories’ dimensionality, looking to identify possible groupings in the five temperament categories evaluated. This technique allows us to represent graphically the relationships among the deer and the behavioral categories. The first two main dimensions were included to group the temperament of the deer before training. The percentage of variance that was explained by these dimensions was 34.3% and 24.2%, respectively. Deer positioned to the left of the x-axis and above the y-axis were classified as less reactive, while the more reactive deer were found to the right of the x-axis and below the y-axis.

A dissimilarity matrix was calculated using the ideal Gower metric for categorical variables to identify temperament patterns. A hierarchical clustering analysis was performed using Ward’s method to minimize variance within groups, and three clusters were defined based on the dendrogram shown in Fig 2.

**Fig 2.**
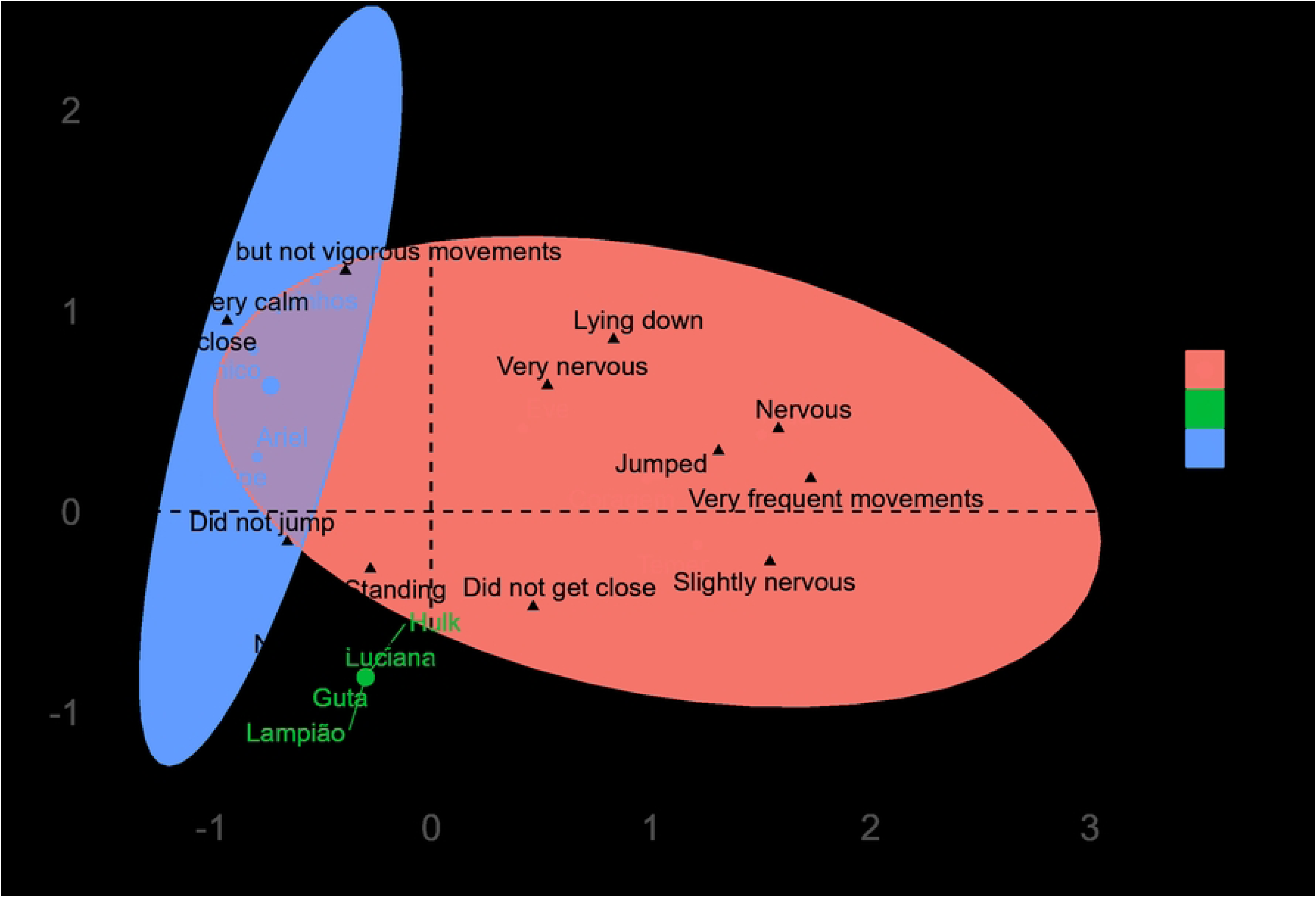
Dendrogram of hierarchical cluster analysis of red brocket deer based on pre-training reactivity categories.

To evaluate the relationship between the coordinates of each deer obtained by the MCA and the number of sessions, the coordinates were treated as a predictor variable in a regression analysis. The correlation analysis showed a strong positive correlation between the coordinates in the first dimension (CD1) and the number of sessions required for habituation (0.724). The coordinates of the second dimension (CD2) did not show significant correlations. Based on this result, the coordinates of the first dimension were included as a predictor variable in a linear regression model. The linear regression model showed how the dimensions obtained in the MCA are related to the number of sessions necessary to carry out the conditioning. The adjusted equation of the model is:

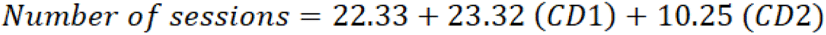

## 3. Results

Fig 3 illustrates the distribution of deer within the formed clusters, highlighting the temperament categories associated with each group before conditioning. It is possible to observe that deer are located close to their temperament classifications, which aids in better understanding the differences among them.

**Fig 3.**
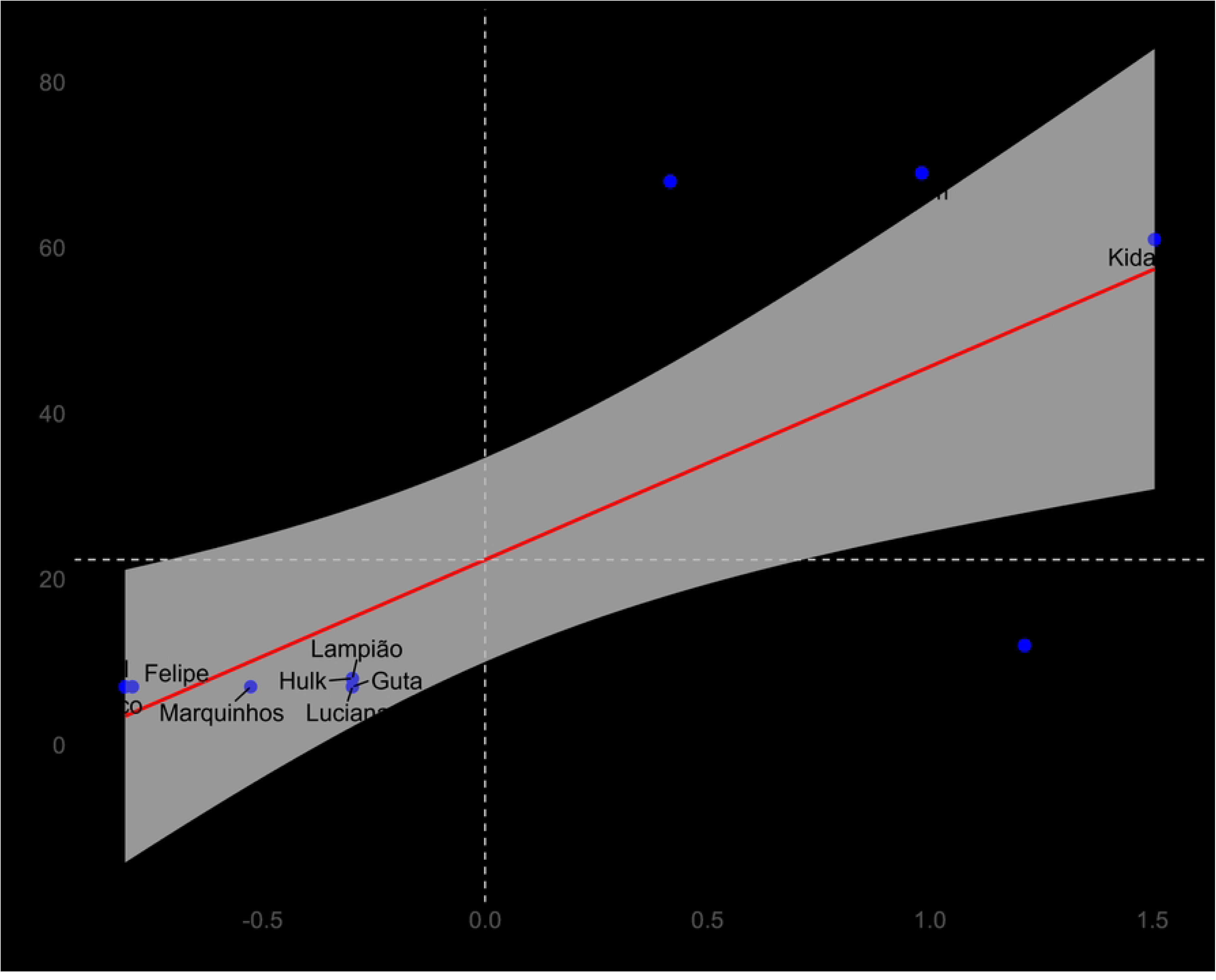
Distribution of deer within the clusters formed and the reactivity categories most attributed or related to the group before starting the training process.

Fig 3 shows that four deer (Kida, Coragem, Eve and Temer), grouped in cluster 1, are more distanced from the others and associated with more negative connotations. A second cluster, composed of Ariel, Chico, Marquinhos and Felipe and a third by Luciana, Guta, Hulk and Lampião, which are associated with positive temperament traits.

The coordinates of the first dimension can be a significant predictor of the regression model, having a greater impact on the number of sessions required for conditioning compared to the coordinates of the second dimension (p-values 0.0076 vs 0.238, respectively). An R2 of 0.596 indicates a moderate fit, suggesting that other factors besides temperament assessment influence the number of conditioning sessions.

The point distribution analysis indicates a linear regression, where the indices of dimension 1 (x-axis) corresponding to the less reactive deer (Ariel, Chico, Felipe, Guta, Hulk, Lampião, Luciana, and Marquinhos) represented by negative values in the MCA (S1 Table). These findings imply that these individuals required fewer habituation sessions than the more reactive deer (Kida, Coragem, Eve), shown separately in Fig 4. The index in dimension 2 did not show a significant relationship with the number of habituation sessions for the trainer.

**Fig 4.**
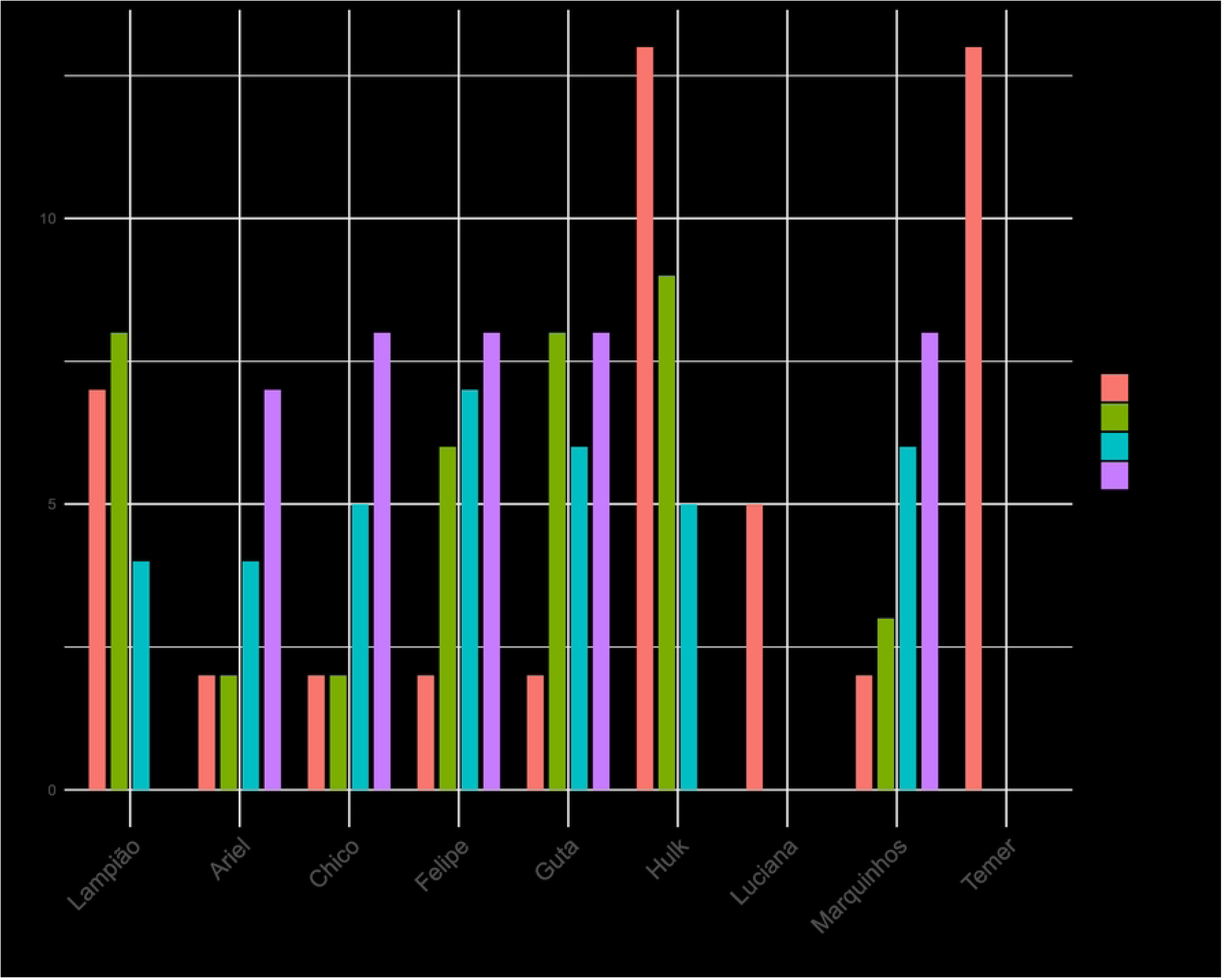
Relationship between the coordinates first dimension of the Multiple Correspondence Analysis (Dimension 1) before starting training sessions and the number of sessions required for habituation to the trainer’s presence in twelve *M. americana*.

After the habituation phase, 9 out of the 12 deer were chosen to undergo training for the commands: approach, snout, belly, and *spray*. Fig 5 highlights the individual progress of each deer for each command.

**Fig 5.**
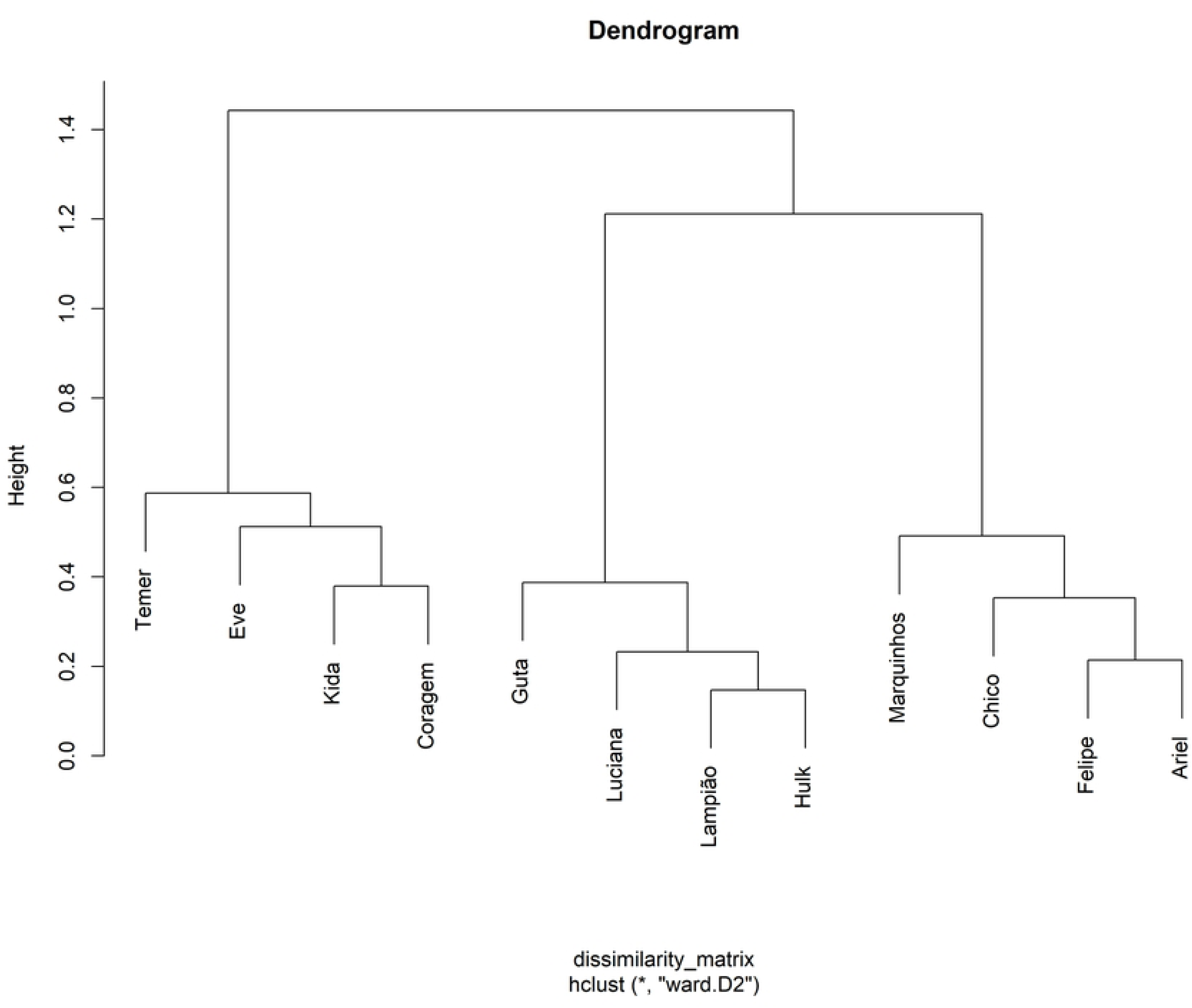
Number of conditioning sessions required for each red brocket deer (n = 9) to learn the commands for veterinary procedures. The bars indicate the commands learned.

## 4. Discussion

Understanding each animal’s personality is essential for improving conditioning techniques [21] and, as a result, increasing training success rates [22]. Results from previous studies show that bolder deer tend to utilize anthropomorphized habitats more efficiently [23] and migrate less due to easier habituation [8]. This relationship suggests that boldness is associated with greater familiarity and ease of adaptation to the environment, thus facilitating learning processes.

The results of this study indicate the need to tailor training strategies to each individual by recognizing the diversity of personalities among deer. For example, when the trainers know the degree of each deer’s reactivity, they can adjust the manner of entering the enclosure and getting closer to the animal, besides defining how to move around it, promoting a more effective and positive connection with the trainer.

Four deer (Ariel, Chico, Felipe, and Marquinhos) habituated to the presence of the trainer in just seven sessions, demonstrating a strong association with positive behavioral categories such as tameness, and no to little movement in the handling box, as assessed before the conditioning process (see cluster 3 in Fig 3). Among the nine deer that participated in the conditioning process, only the four mentioned, along with Guta, were able to learn the command “*spray*,” representing a success rate of 41.66% in learning all trained commands. Animals exhibiting greater curiosity and exploratory behavior, associated with positive behavioral traits, generally show better adaptation to stressful situations, a greater willingness to participate in training [24], and improved behavioral flexibility [25]. Similar responses were observed in *Cervus canadensis*, where bolder individuals were more responsive and learned more quickly [7].

Nervous animals tend to adapt poorly to changes [1,11]. This study yielded results consistent with these observations, indicating that more reactive and fearful deer faced significant difficulties in habituating to the trainer’s presence and learning new commands. In other words, the progression of these deer throughout the study was slower [26].

Three deer (Kida, Eve, and Coragem) exhibited the worst temperament, which persisted throughout the initial stage of the study, undergoing more than 60 sessions of habituation each, as shown in Fig 4, which illustrates that those with the most negative indices tend to take longer to habituate. This trend is also observable in cluster 1 of Fig 4, where they remain close to characteristics categorized as slightly nervous, nervous, or very nervous animals and do not get close to the unfamiliar person, corroborating the results of Huang et al. [27]. These deer attempted to jump during the handling procedures reactivity test and exhibited more intense movement when kept in the handling box. This behavior aligns with the characteristics commonly attributed to the shyness of forest musk deer (*Moschus berezovskii*) [28] and alpine musk deer (*Moschus chrysogaster*) [10]. Similar results were found in horses [29], showing that the poorest learning performances were observed in animals displaying negative behaviors.

The other four deer (Guta, Hulk, Lampião, Luciana) showcased distinct individualities linked to their personalities. Fig 5 illustrates that not all these deer mastered every command except for Guta. There is a correlation with their temperament; calmer individuals learned three or four commands, while the others learned only one or two.

It should be highlighted that each deer learned at its own pace, demonstrating significant individual variation in the learning process due to their distinct temperaments. This finding aligns with other studies suggesting that differences in responses to temperament tests are influenced by the unique personalities of the animals [7]. Furthermore, this supports the notion that personality is directly related to success in operant conditioning [30].

## 5. Conclusions

This study revealed that less reactive and calmer individuals had a greater learning capacity and accurately executed most of the commands applied, while more reactive and fearful individuals exhibited a slower progression in mastering those commands. These findings underscore the significance of temperament in the learning process of deer, suggesting that individual differences play a vital role in how animals respond to training. Recognizing these differences can inform more effective training strategies and improve overall outcomes in efforts to condition wildlife.

## Acknowledgements

We express our sincere gratitude to the coordinator (Prof. José Mauricio Barbanti Duarte) and staff of the Deer Research and Conservation Center (NUPECCE) at the Faculty of Agricultural and Veterinary Sciences, UNESP, Jaboticabal campus, SP, Brazil, for their invaluable support and assistance throughout this study. We also appreciated the support of the Graduate Program in Animal Science at São Paulo State University, Faculty of Agricultural and Veterinary Sciences, Jaboticabal campus. This study was financed in part by the Coordenação de Aperfeiçoamento de Pessoal de Nível Superior -Brasil (CAPES) – Finance Code 001.

## CRediT authorship contribution statement

**Lara Caveanha Gragnanello**: Conceptualization, Data curation, Funding Acquisition Investigation, Methodology, Project administration, Resources, Validation, Visualization, Writing – original draft, Writing – review & editing, **Mariana Parra Cerezo:** Data curation, Formal analysis, Investigation, Validation, Writing – review & editing, **Cristiane Schilbach Pizzutto:** Conceptualization, Data curation, Methodology, Visualization, Investigation, Writing – original draft, Writing – review & editing and **Mateus J. R. Paranhos da Costa**: Conceptualization, Data curation, Formal analysis, Methodology, Supervision, Visualization, Writing – original draft, Writing – review & editing.

